# A Specific Ancestral Sequence Reconstruction of Flagellin Can Drive Motility in *Salmonella enterica* but not *Escherichia coli*

**DOI:** 10.64898/2026.10.20.760431

**Authors:** Jacob Scadden, Jamiema Sara Philip, Pietro Ridone, Yoshiyuki Sowa, Matthew A.B. Baker

## Abstract

The bacterial flagellar filament, built from thousands of flagellin subunits, is essential for flagellar motility. Here, we aimed to use ancestral sequence reconstruction (ASR) to generate conservative, ancestral flagellin sequences that surround the divergency point between *Escherichia* and *Salmonella* species and tested their ability to restore motility in flagellin deleted *Escherichia coli* and *Salmonella* Typhimurium. Only one (N131) out of three candidates restored motility in *S.* Typhimurium but not *E. coli*, despite high structural similarity to native FliC/FljB. *S.* Typhimurium expressing N131 showed reduced motility relative to wild-type *Salmonella* FliC, alongside increased tumbling frequency. Serial passage across swim plates produced a derivative, N131.10, containing a single flagellin mutation (M144I), with a larger swim ring diameter but also increased tumbling frequency. Comparison of structural interaction networks among the residues of *S.* Typhimurium LT2 FliC with those of other filament proteins showed that non-motile candidates (N132 and N134) carried substitutions that disrupted key core-domain atomic interactions. These results demonstrate that flagellar filament assembly tolerates only limited sequence variation and that this capacity for tolerance differs between *E. coli* and *S.* Typhimurium despite their close phylogenetic relationship.

**Importance:** Bacteria swim using long, rotating filaments assembled from thousands of copies of a protein called flagellin. We reconstructed likely ancestral versions of this protein to investigate how evolutionary changes affect swimming. Of three versions tested, only one restored swimming in *Salmonella* unable to make its own flagellin. The same protein did not work in the closely related bacterium *E. coli*. After repeated selection for better movement, a single additional change helped the *Salmonella* spread farther and also altered its swimming behaviour. These findings show that closely related bacteria differ in the changes that their flagellar assembly machinery can tolerate. Understanding these limits helps explain how flagella evolve while retaining their function and provides a foundation for studying the role of flagellar variation in immune recognition and evasion.

## Introduction

The bacterial flagellar motor (BFM) is a nanomachine that enables bacterial cells to swim (1). Flagellar-driven motility is an ancient trait and is found in multiple bacterial species across the bacterial tree of life (2,3). Approximately 30 core structural genes form the BFM in *Escherichia coli* and *Salmonella enterica* subsp. *enterica* serovar Typhimurium (*S*. Typhimurium), which form the basal body, stator, hook and filament sub-structures (1,4). The filament acts as the propeller of the system, rotating to generate thrust and driving the bacterium through its environment (5). The flagellar filament is primarily made of a protein called flagellin and its presence in the genome is critical for motility (3,6). Bacterial flagellar filaments are made of approximately 20,000 flagellin subunits that arrange themselves in a 11-start helix that supercoil to form a helical superstructure that generates thrust for the bacterium (7,8).

There is a wide diversity of flagellins found across the bacterial kingdom (9). *E. coli* K-12 is monophasic with a single copy of flagellin which contains two conserved core domains (D0 and D1) and two hypervariable outer domains (D2 and D3), however these outer domains are not always present in other bacteria (5,8,10,11). Some bacteria, such as *S*. Typhimurium, contain multiple copies of flagellin (FliC and FljB) which can undergo changes in expression (phase variation) to allow for motility in complex niches and allow for bacterial escape in phage-rich or high host immune response environments (12). Conversely, other *Salmonella* species, such as *S*. Typhi, have a monophasic flagellin gene cluster similar to that of *E. coli* (13,14).

*Escherichia* diverged from *Salmonella* approximately 100-160 million years ago and it is hypothesised that some *Salmonella* species gained or duplicated an additional flagellin gene to create the biphasic system present today in some species (15,16,13). There is evidence of interchangeability of flagellins between these genera, with previous work demonstrating that flagellins from the same family (*E. coli* flagellin expressed in *Salmonella*) as well as chimeric flagellins can form filaments (17–21). However, by generating ancestral flagellin candidates surrounding the divergence point between *E. coli* and *Salmonella*, we can identify the critical residues required for filament formation.

Ancestral sequence reconstruction (ASR) is an *in silico* approach that uses alignments of extant sequences and their phylogenetic relationships to estimate ancestral sequences at internal nodes of the phylogeny (22,23). Previous work has generated ASR constructs of the MotA and MotB stator unit proteins which were then screened for functionality in stator deleted *E. coli*, resulting in rescue of motility for some candidates (24,25). Here we utilised this technique to generate ancestral flagellin candidates that represent common ancestors surrounding the divergence point between *E. coli* and *Salmonella* sp. that were cloned and expressed in Δ*fliC* and Δ*fliC* Δ*fljB E. coli* and *S*. Typhimurium strains, respectively. The resulting motile flagellin candidate demonstrates that ASR can generate motile filaments that can be further optimised through subsequent mutation after serial passages.

## Materials and Methods

### Bacterial strains, plasmids and growth conditions

All bacterial strains and plasmids used in this study are described in Table S1. The *E. coli* SHU649 (Δ*fliC*) and *S*. Typhimurium TM113 (Δ*fliC* Δ*fljB)* (26) mutants were generated using Lambda (λ) RED recombineering, as previously described (27), to delete the *fliC* and the *fliC* and *fljB* genes, respectively. Both bacterial species were grown in lysogeny broth (LB) (10 g/L NaCl, 10 g/L Bacto tryptone, and 5 g/L yeast extract) and tryptone broth (TB) (5 g/L NaCl, 10 g/L Bacto tryptone). To make solid LB agar (2%, wt/vol) and LB swim agar (0.3%, 0.35% and 0.4% wt/vol) plates 20.0, 3.0, 3.5, or 4.0 g/L agar was added, respectively. Bacterial cultures were all grown at 37℃, unless for swim plates and growth in TB for free swimming assays which were performed at 30℃. Antibiotic selection used ampicillin (AMP) at 100 µg/mL. Plasmid gene expression induction used 0.1 mM isopropyl β-D-1-thiogalactopyranoside (IPTG).

### Generation and selection of flagellin ASR candidates

To generate flagellin ASR candidates, first, representative flagellin amino acid sequences were downloaded from UniProt (28). The term “flagellin” was used in the UniProtKB database followed by sequence refinement by searching against the Taxonomy filter for “Bacteria – [2]” and a size exclusion parameter of 490-510, identifying 3047 flagellin sequences. These sequences underwent UniRef90 ID mapping to remove similar sequences. These sequences were then manually assessed to remove any fragmented sequences and ensuring a coverage across the GTDB bacterial tree of life (29). This resulted in 78 representative bacterial flagellin amino acid sequences being selected. Representative flagellin sequences were aligned using MAFFT (30), viewed on AliView (31), followed by generation of a phylogenetic tree using IQTree on the CIPRES Science Gateway (32) and visualisation using iToL v 7.6 (33).

Generation of ASRs from both the bacterial and Gammaproteobacteria alignment and phylogenetic tree using CODEML in PAML (v4.10.3) (34) under the JTT amino acid substitution model. Among-site rate variation was modelled using discrete gamma rate categories (ncatG = 8), with the gamma shape parameter estimated (fix_alpha = 0). Three nodes were selected, N131, N132 and N134 based on the proximity to the extant sequence of *E. coli* K-12 FliC. Raw ASR candidate sequences were aligned using MAFFT (30) with both *E. coli* and *S*. Typhimurium FliC (P04949 and P06179, respectively) and additional residues in the ASR candidates were removed in conjunction with the probability score of each residue (Table S2). Amino acid sequences of curated ASRs were reverse transcribed in Benchling and manually codon optimised against the *E. coli* K-12 *fliC* nucleotide sequence (GenBank: M14358.1).

### Flagellin structural prediction and outer domain structural similarity

AlphaFold3 was used to predict the structure of 78 representative flagellin sequences and the three candidate flagellin ASRs (35). Predicted structures were visualised with using PyMol (v3.1.6.1). Outer-domain (D2/D3) similarity among 78 bacterial flagellins was assessed in parallel by sequence and by structure, following the general scheme of Fields et al., (9). AlphaFold model_0 predictions were parsed with Biopython’s MMCIFParser to recover Cα coordinates in residue order. Outer domains were isolated by fixed-index trimming of the first 163 and last 100 Cα positions, which removes the terminal D0/D1 helices; models retaining fewer than 10 residues after trimming were discarded. A ClustalW-format multiple alignment of the full-length flagellins was read with Bio.AlignIO. Alignment columns were mapped onto *Escherichia coli* FliC (P04949) residue numbering, and columns corresponding to P04949 residues 181–400 were taken as the outer-domain block. Pairwise identity was computed over this block as the fraction of identical positions among columns ungapped in both sequences. For each pair differing in outer-domain length by <30%, the shorter chain was compared against a centred, equal-length contiguous slice of the longer chain, superposed by Kabsch rotation (SVD), and scored as mean 1/(1 + (dᵢ/d₀)²) with d₀ = 1.24(L − 15)^(1/3) − 1.8, L = max length. Pairs failing the length criterion were scored 0. This sequence-order-dependent, single-slice approximation performs no alignment search, so scores are lower bounds on true TM-score. Matrices were cached to JSON. Three undirected graphs were built with NetworkX over all 78 nodes: sequence identity >0.20, TM-score >0.5, and combined max (TM-score, identity) >0.45, with similarity as edge weight. Communities were detected by weighted greedy modularity maximisation (Clauset–Newman–Moore). Layouts used spring (Fruchterman– Reingold) embedding of the connected subgraph with a fixed seed, singletons arranged on a surrounding circle, nodes coloured by bacterial class. Community assignments and all three matrices were exported as CSV/JSON.

### FliC residue interaction analysis with other filament proteins

Interchain residue contact networks for Salmonella flagellin were determined using the RING 3.0 server (Residue Interaction Network Generator; https://ring.biocomputingup.it/) applied to two cryo-EM structures: PDB 9GNZ (flagellar cap-filament complex; 22 chains comprising FliC subunits A–Q and FliD subunits R–V) and PDB 9GO6 (hook-filament junction complex; 50 chains comprising FliC subunits a–o, FlgK subunits A–K, FlgL subunits L–V, and FlgE subunits 1–9 and w–z). Calculations were performed under strict distance thresholds with all interaction types enabled (hydrogen bonds, van der Waals contacts, π–π stacking, and ionic interactions). The resulting edge files (TSV format; NodeId notation Chain:ResNum:_:AminoAcid) were parsed to retain only interchain contacts, with intrachain interactions excluded. For each structure, contact partners of all FliC chains were pooled and mapped to residue positions in the UniProt P06179 reference sequence. Positions were classified by partner identity (FliC–FliC, FliC–FliD, FliC–FlgL, FliC–FlgK, FliC–FlgE) with positions contacting multiple partner types flagged accordingly. Structural domain boundaries followed UniProt P06179 annotation (D0·N: 1–67; D1·N: 68–175; D2·N: 176–237; D3: 238– 273; D2·C: 274–314; D1·C: 315–400; D0·C: 401–495). Combined contact data from both structures were visualised on the 9GNZ chain A monomer in PyMOL 3.0.

### Cloning of ASR candidates into expression vectors

All ASR candidates were produced as DNA fragments by TWIST Bioscience, Decode and were amplified using ASR_F and ASR_R (Table S3), followed by PCR product purification (Qiagen). were cloned into the pTrcHis_C (ThermoFisher Scientific) using restriction digestion (NcoI-HF and EcoRI-HF; New England Bioscience (NEB)), followed by ligation using T4 ligase (NEB) and transformation into *E. coli* 10-β (NEB), as per manufacturer’s instructions. Correctly cloned vectors were identified using colony PCR using pTrc_F and pTrc_R (Table S3) purified using Qiagen Miniprep kit and confirmed via Sanger sequencing (Ramaciotti Centre for Genomics). Plasmids were transformed by electroporation into *E. coli* SHU649 and *S*. Typhimurium TM113. Transformants were selected on ampicillin (100 µg/mL) agar plates.

### Swim Plate Assays

Swim plate assays were performed as previously described (36). Briefly, 0.3% agar plates were made with appropriate antibiotics and inducers, with 1 µL of induced overnight culture used to spot the plates. Plates were incubated for 16 hrs at 30℃. A Bio-Rad Gel Doc XR was used to image plates. Swim ring diameters were analysed using ImageJ (FIJI) (37) and data were analysed and displayed using GraphPad Prism 10 (v10.5.0).

### Free-swimming Assays

FliC-ASR expressing *S*. Typhimurium and *E. coli* SHU649 overnight cultures were grown at 37°C, shaking (180 rpm). A fresh culture was made using 5 mL TB with appropriate antibiotics and inducers. When the cultures reached OD_600_ 0.2, 5 µL of culture was flowed into a glass tunnel slide, as described in (38). Prior to inoculation of the tunnel slide 5 µL of Bovine Serum Albumin (BSA) (100 µg/mL) was added to prevent adhesion of bacteria to the glass slide and cover slip. Videos were recorded using an Evos M7000 at 30 frames per second for 13 seconds (∼400 frames) and analysed using cell tracking software on Labview 2019 software (National Instruments). Individual cell speeds, tumble frequency and tumble bias were recorded and were analysed and displayed using GraphPad Prism 10 (v11.9.2).

### Flagellar filament staining

Bacterial species were grown overnight in LB broth at 37℃ in a shaking (180 rpm) incubator, containing all appropriate antibiotics and inducers. On a 0.3 % LB agar plate, 1 µL of culture was used to inoculate the medium, containing all appropriate antibiotics and inducers and incubated at 30°C, static overnight. A drop (∼10 µL) of ultrapure water was spotted on the edge of the swim ring and allowed to sit for 2 mins. A sterile inoculation loop transferred the drop to a glass slide mixing with a drop of ultrapure water. A cover slip was added, followed by 8 µL of Remel Flagella stain (Thermo Fisher) and incubated at room temperature for 10 mins. The cells were imaged using a Leica DM750 microscope at ×100 magnification and captured using a MEE-500 camera (Yegren, China) and the camera app on Windows 10 (v.10.0.1.19045). The lengths of the filaments were measured manually on ImageJ (FIJI). Data were analysed and visualised using GraphPad Prism 2.0 (v11.9.2).

### Growth Curve Assays

Overnight cultures were grown in LB broth with appropriate antibiotic and inducer concentrations at 37°C, shaking. The following day, a 1:100 dilution was made in LB media containing appropriate antibiotic and inducer concentrations. To a 96-well plate 99 µL LB broth was added followed by 1 µL of the 1:100 dilution of the overnight culture. Six replicates were used for each strain. The OD_600_ was measured every 20 mins using a POLARstar Omega and incubated for 16 hrs at 37°C. Data were blank corrected and analysed and visualised using GraphPad Prism 2.0 (v11.9.2).

### Whole genome sequencing of N131 variant evolved *S.* Typhimurium

To identify chromosomal mutations in N131 variant expressing *S*. Typhimurium TM113, six variants were sent for whole genome sequencing at the Ramacciotti Centre for Genomics using a MiSeq 2 × 150–base pair (bp) chip on an Illumina sequencing platform. Single nucleotide polymorphisms were identified using Snippy and aligned to the *S*. Typhimurium LT2 genome (GenBank: AE006468.2) (39).

### Supernatant protein precipitation and SDS-Page

Bacterial strains containing ASR plasmids were cultured from glycerol stocks in 10 mL LB media overnight then sub-cultured in 50 ml pre-warmed media with AMP addition at 30°C. At OD_600_ = 0.4, cultures were induced with 100 mM IPTG and incubated for another 3 hrs at 30°C. Cultures were heated to 60℃ for 10 mins. The cells were harvested by cell centrifugation (4000 x g, 15 mins, 4°C). Culture supernatants collected by centrifugation were filtered with a 0.22 μm filter. Supernatant total proteins were precipitated and concentrated from the filtered supernatant by addition of 100% cold TCA to give 20 % and incubated at 4°C for 2 h or overnight. Concentrated and precipitated proteins were pelleted by centrifugation (4000 x g, 30 mins, 4°C), washed with 1/5th volume ice cold acetone for 2 hrs at -20°C, recentrifuged, supernatant removed then pellet dried at room temperature until acetone was completely evaporated. Pellet was stored at -20°C and resuspended in 50 mM sodium acetate (pH 5.5) until fully dissolved. Protein quantification was performed using Bradford’s reagent as described (40). A NuPage 4-12% Bis-tris SDS-Page gel was used to visualise protein content as per manufacturer’s instructions Briefly, 10 µg of crude supernatant protein was loaded with LDS and reducing agent, heated for 10 mins at 70℃ and ran at 200V for 25 mins, stained using BlueSafe and destained overnight in dH_2_O. Gels were imaged using a Bio-rad GelDoc Transilluminator.

## Results

### Flagellin ASR candidates share conservation of core domains but have varied outer domains

The phylogenetic tree was generated using 78 representative flagellin sequences with lengths ranging from 490-510 amino acids from major taxonomic classes of flagellated bacteria (Alphaproteobacteria = 10, Betaproteobacteria = 10, Gammaproteobacteria = 37, Epsilonproteobacteria = 6, Clostridia = 5, Chthonomonadia = 1, Syntrophia = 1, Terriglobia = 1, Aquificia = 1, Actinomycetes = 1, Bacilli = 1, Phycisphaerae = 2, Unclassified = 2), showed clustering primarily based on the taxamonic class (Figure 1A). We decided to use amino acid sequence length as an exclusion criterion to minimise the gaps within the alignment and aid in the ASR output sequencing trimming due to the high variability of the outer domains of flagellin and that the wild-type flagellins from our *E. coli* and *S*. Typhimurium expression systems are within this range. Analysis of predicted outer domain structures showed ∼46% (36/78) of the representative flagellins clustered in four main groups based on structural homology (Figure 1A and S1A and B).

**Figure 1.**
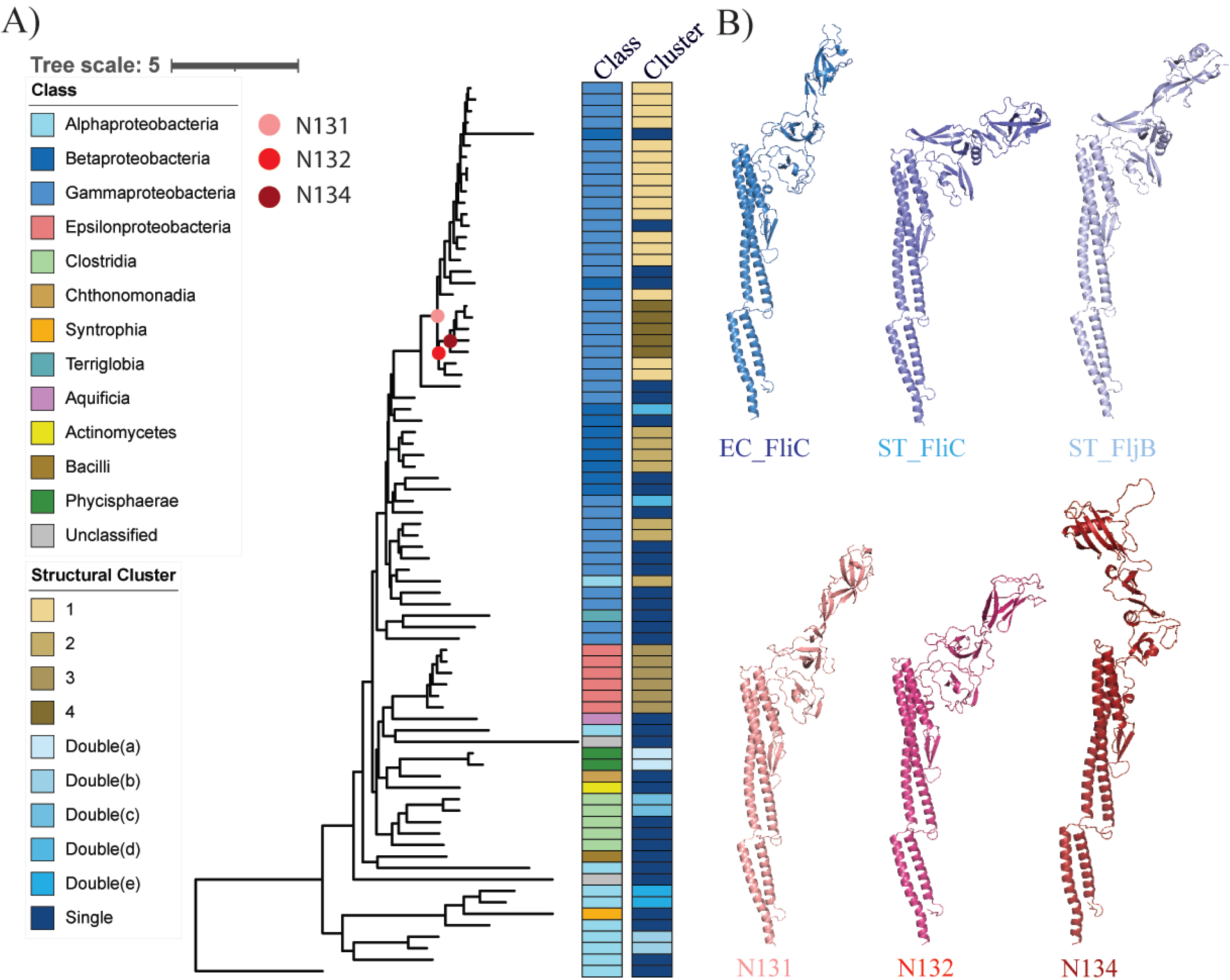
A) Phylogeny of 78 representative flagellin amino acid sequences from across the bacterial tree of life. Inner column colours represent Class. Outer column represents the outer domain structural cluster to which the predicted structural model belongs. The three nodes selected for further study are indicated on the phylogeny by coloured dots; Light red = N131, Red = N132 and Dark red = N134. B) Alphafold 3 predicted models for the tertiary structures of FliC (*E. coli* K-12 (EC)), FliC and FljB (*S*. Typhimurium LT2 (ST)) and highest confidence AlphaFold3 models of N131, N132 and N134.

The percentage identity of the whole flagellin amino acid sequences ranged from 15.73% to 64.17% compared with *E. coli* K-12 FliC (Figure S2A). As expected, MAFFT alignment showed higher conservation in the N- and C-termini (82.64% to 15.87%, mean = 50.92% ± 20.3) than in the outer, variable domains (27.18% to 10.50%, mean = 16.70% ± 4.38) (Figure S2B and C), when compared to FliC from *E. coli* K-12 (P04949). Three ASR candidates, N131, N132 and N134, were selected based on their proximity to the extant node of *E. coli* K-12 FliC (EC_FliC) (Figure 1A). Structural homology analysis of the extant leaves after each node of interest were members of clusters 1 (n=17), 4 (n=5) and single structural morphologies (n=4) (Figure 1A, S1A and B, S4). Assessment of probability score of each residue after manual curation of the sequence of *E. coli* K-12 FliC showed that lowest confidence residues are found between residues ∼175 to 420 (Figure S3A-F and S4A-C). N131, N132 and N134 all had over half their residue probabilities ≥0.9 at 61.8%, 62.3% and 58.2%, respectively (Figure S4D-F, Table S2). The ASRs produced were upstream of only Gammaproteobacteria and Betaproteobacteria, predominantly *Salmonella*, *Citrobacter* and *Escherichia* species (Figure 1A and S5). All ASR candidates had the FliS export recognition sequence maintained between positions 26 to 47, except position 28 where the serine in EC_FliC was replaced in all ASR candidates by an alanine (Figure S6). The alanine, however, is conserved in both *S*. Typhimurium LT2 FliC (ST_FliC) and FljB (ST_FljB) (Figure S6).

### N131 rescues motility in *S*. Typhimurium but not *E. coli*

All ASRs were initially tested in *E. coli* SHU649, where no motility was observed after 16 hrs and 48 hrs (Figure 2A and B, Figure S7). As all the ASRs were selected from nodes preceding extant *Salmonella* species (Figure 1A), we tested each ASR in *S*. Typhimurium TM113. We found that a single ASR candidate, N131, was able to rescue motility in this bacterial expression system (Figure 2C and D). The diameter of the swim ring was significantly reduced in N131 (12.9 ± 0.9 mm) compared to wild-type FliC (60.3 ± 3.1 mm) (Figure 2D). However, the swim ring diameter was significantly greater than the non-motile growth of N132, N134 and the negative pTrc control (Figure 2D). Growth curves for all expression species showed final OD_600_ values of between 0.854 to 1.00, with *S*. Typhimurium pTrc having the fastest growth rate (Figure S8).

**Figure 2.**
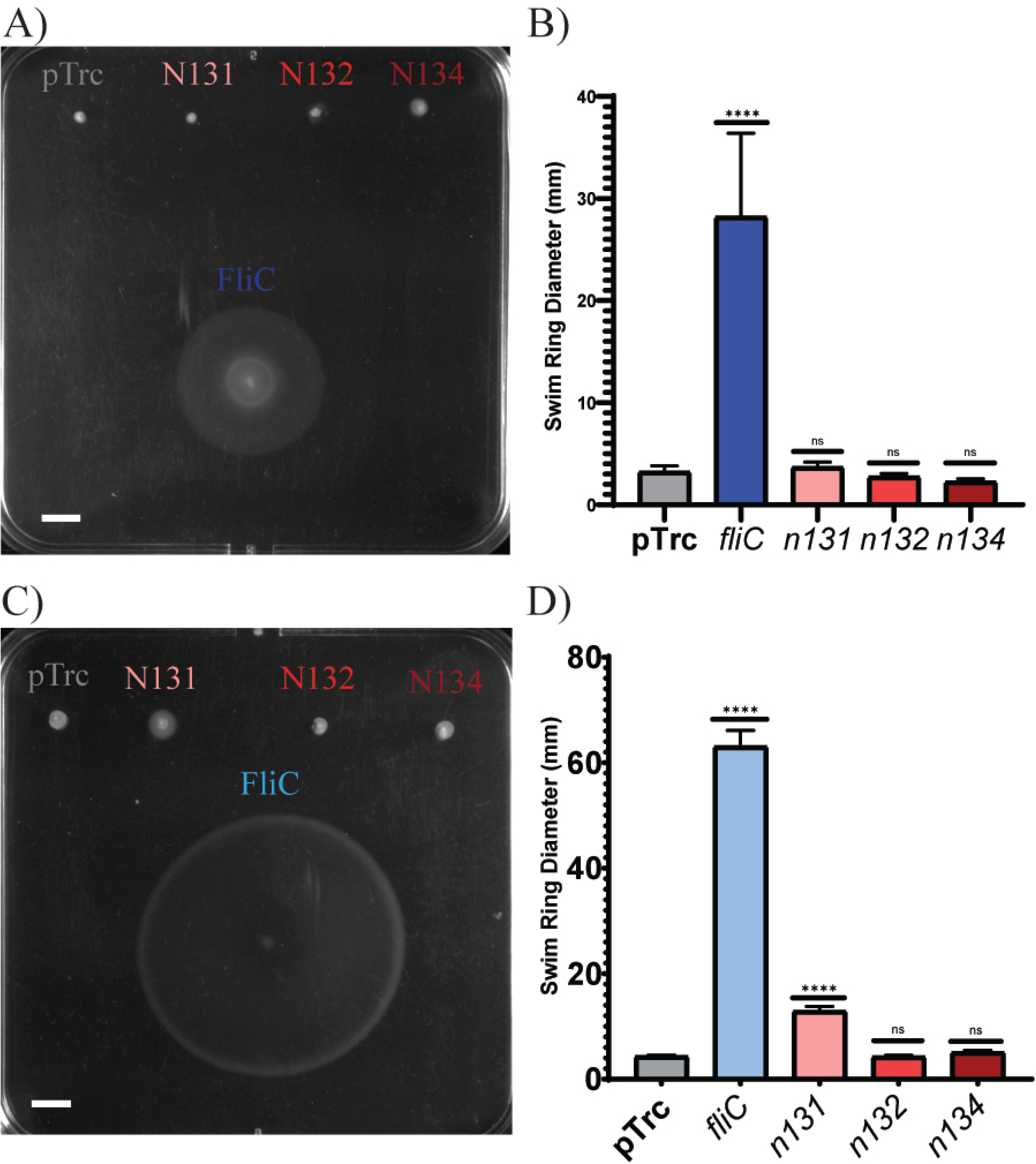
Representative swim plates (0.3% agar) for A) *E. coli* SHU649 expressing wild type *fliC* and ASR candidates with B) corresponding swim ring diameters quantified. C) An example swim plate for *S*. Typhimurium TM113 expressing wild type *fliC* and ASR candidates with D) swim ring diameters quantified. An empty vector (pTrc) was used as a negative control in both expression strains. In all swim ring diameter graphs the error bars represent standard deviation. Statistical significance against the negative pTrc control was determined using a one-way ANOVA (P > 0.05 = ns, P ≤ 0.0001 = ****). White bars represent 10 mm.

Staining of the flagellar filaments from wild-type FliC and N131 did not show a significant decrease in filament length (3.6 ± 1.3 µm to 2.9 ± 0.8 µm for wild-type FliC and N131, respectively) or number of visible flagella per cell (1.7 ± 0.9 and 1.6 ± 0.8 for wild-type FliC and N131, respectively) (Figure 3A-C). Free swimming analysis observed a ∼4-fold decrease (14.2 µm/s to 4.3 µm/s) in swim speed compared to wild-type FliC (Figure 3C). Conversely, tumble frequency was significantly increased in *n131* expressing *S*. Typhimurium TM113 (Figure 3E). Precipitated protein from the heat-treated supernatants of the ASR cultures showed that N131 and N132 were present in the supernatants of both *E. coli* SHU649 and *S*. Typhimurium TM113 with no clear band observed for N134 (Figure S9).

**Figure 3.**
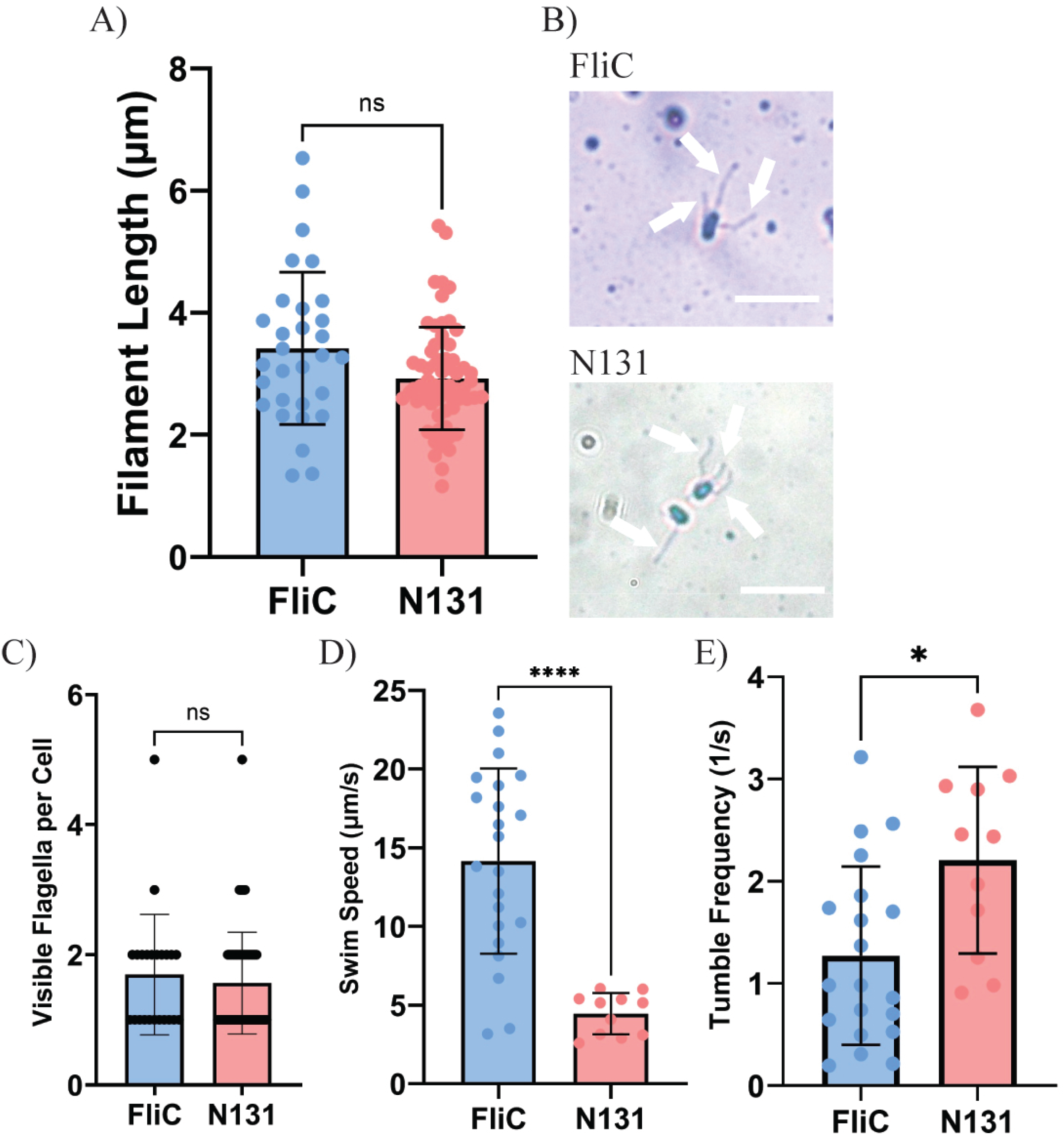
A) Filament lengths for wild type *fliC* and *n131* expressing *S*. Typhimurium TM113. B) Representative images of Remel stained flagellar filaments formed by FliC and N131. White bars represent 5 µm. White arrows point towards stained flagella. C) Mean visible flagella number per cell, D) bacterial cell swim speeds and E) tumble frequency of motile *fliC* and *n131* expressing *S*. Typhimurium TM113. In all graphs, error bars represent standard deviation. A Welch’s t-test was used to assess significance (P > 0.05 = ns, P ≤ 0.05 = *, P ≤ 0.0001 = ****).

### Evolution of N131 in response to sequential passaging and increasing viscosity

Following the motility phenotype seen in the *n131* expressing *S*. Typhimurium TM113, this variant was subsequently passaged nine times to observe if there would be any improvement in motility. When compared to the parent (N131.1) swim ring diameter the final variant (N131.10) had a 2.2-fold increase in swim ring diameter (Figure 4A and B). Examination of growth curves for each variant indicated that the growth rates for all passaged N131 variants were the same (Figure S10). Sanger sequencing analysis of the *n131* gene indicated a single nucleotide mutation at position 432 after the second passage (N131.2 to N131.3) which causes a change to the methionine at position 144 to an isoleucine (Figure 4C and D, and S10). AlphaFold 3 structural predictions for both N131_M144, N131_I144, *E. coli* K-12 FliC and *S*. Typhimurium LT2 FliC were analysed using RING which showed that, although the I144 maintained the hydrogen bond to I158 of the beta-loop, there was no predicted additional interaction to the V135 (N131_M144 and *E. coli* FliC) or I74 (*S*. Typhimurium) (Figure 4C). No further mutations in the *n131* gene were observed in the final passaged variant, N131.10 (Figure S11). There was no significant change in free swimming speed between N131.1 and N131.10 (Figure 4E), however the tumble frequency was significantly higher for N131.10 compared to the parental N131.1 and wild type *S*. Typhimurium FliC (Figure 4F). Whole genome sequencing identified mutations in passaged variants when compared to the parental N131.1 variant (Figure 4G). A mutation in *fliN* (R70H) was observed in all passaged variants sequenced, additionally a mutation in *fliD* (G324R) was seen after variant N131.7 with N131.10, although the latter had a read frequency of ∼56 %) (Figure 4G).

**Figure 4.**
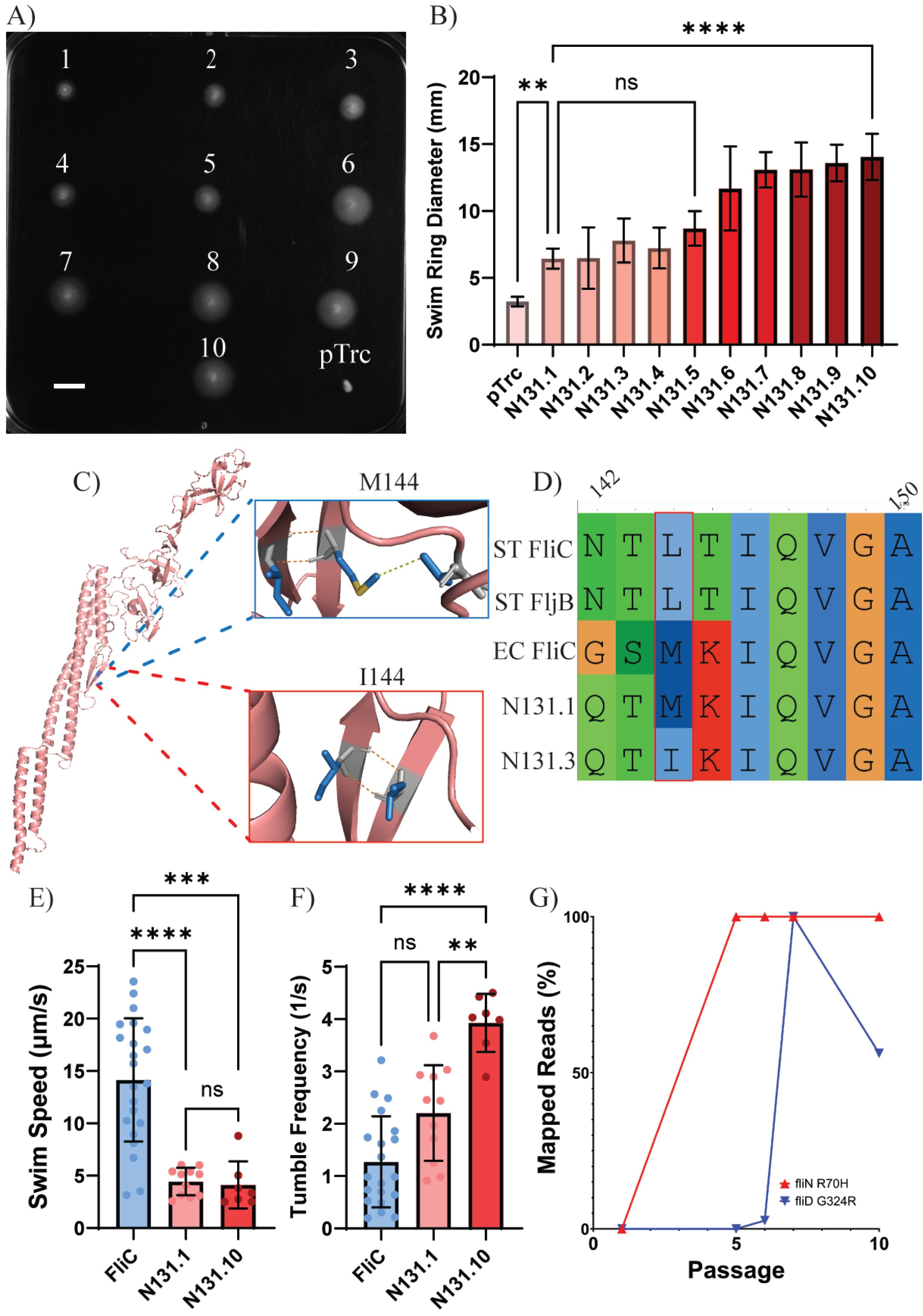
A) Representative swim plate of *S*. Typhimurium TM113 expressing *n131* passaged variants in 0.3% agar. White bar represents 10 mm. B) Swim ring diameters of N131 passaged S. Typhimurium TM113. Error bars represent standard deviation. pTrc was used as a negative control. C) M144I mutation (Blue) mapped on N131 Alphafold 3 predicted structure (Pink). Contact changes with beta-loop structure of D1 domain are shown alongside. H-Bonds (Orange) and Van der Waals (Yellow) interactions are shown. D) Alignment comparing the M144I mutation (Red box) in context with *E. coli* K-12 FliC, *S*. Typhimurium LT2 FliC and FljB and the parent N131 amino acid sequence. E) Average swim speed, F) tumble frequency of ST_FliC, N131.1 and N131.10. Error bars represent standard deviation. Statistical significance against the mean value of each flagellin variant was determined using a two-way ANOVA (P > 0.05 = ns, P ≤ 0.05 = *, P ≤ 0.01 = **, P ≤ 0.001 = ***, P ≤ 0.0001 = ****).

In higher-viscosity agar (0.35% and 0.4%), most passaged N131 variants showed significantly larger swim rings than parental N131.1 (Figure S12A-D). After one week at 30°C on 0.4% agar plates, motile flares from N131.2, N131.4, N131.5 and N131.7 were re-passaged; N131.7.1 showed the greatest increase in swim ring diameter (Figure S13), with no additional *n131* identified via Sanger sequencing of the purified plasmid (Figure S11). However, an additional chromosomal mutation was identified in *motA* (G199V) (Table S4). Furthermore, when N131.1 and N131.10 were re-transformed into *S*. Typhimurium TM113 the latter still producing significantly larger swim rings, although swim speed and tumble frequency was not significantly different (Figure S12E-H).

### N131 contains residues that are key for filament formation

The inter- and intra-subunit interactions of the FliC of *S*. Typhimurium LT2 were analysed using Residue Interaction Network Generator (RING). The cryo-EM structures of the filament- cap (PDB 9GNZ, (41)) and filament-hook complexes (PDB 9GO6, (41)) (Figure 5A and B) were initially analysed to identify the key interacting residues between FliC and other proteins within the filament such as the filament cap (FliD), the hook-filament junction (FlgL/FlgK) and the hook (FlgE). Across the two cryo-EM structures, the 17 FliC subunits, interacted with FliD (59 FliC residues) and FlgL (67 FliC residues), with a single interaction with FlgK (1 FliC residue) residues (Table 1). There were no direct interactions between residues involving FliC and FlgE. There were 157 FliC residues that were able to form contacts with other FliC monomers from *S*. Typhimurium LT2 across the two structures (Table 1).

**Figure 5.**
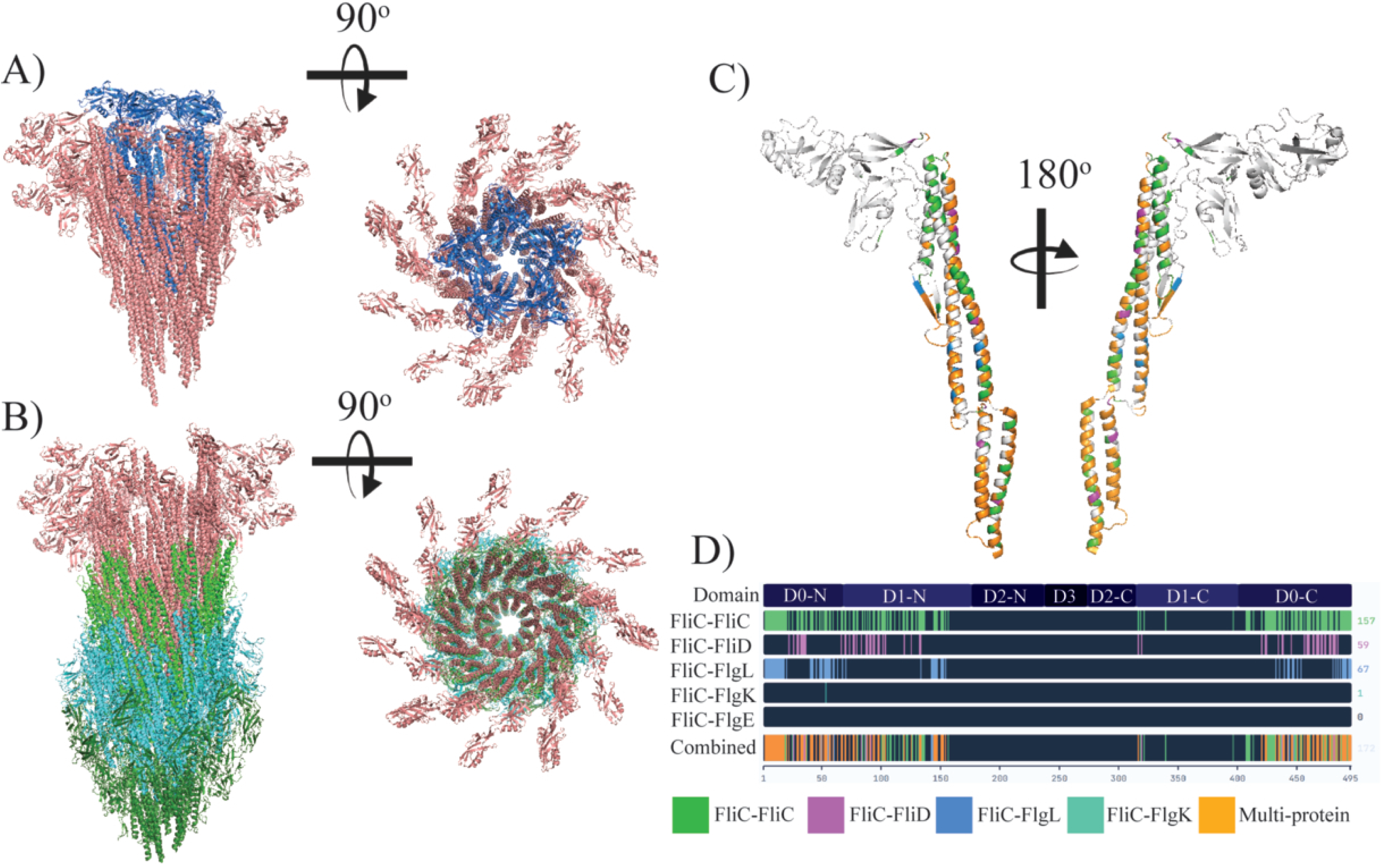
Cryo-EM structure of the A) filament-cap (9GNZ) and B) filament-hook complexes (9GO6) (41). A side view (Left) and top view (Right) show the helical structure of the filament. FliC = Pink, FliD = Blue, FlgL = Green, FlgK = Turquoise, FlgE = Dark Green. C) Cryo-EM model of *S*. Typhimurium LT2 FliC with interacting residues highlighted. FliC-FliC = Green, FliC-FliD = Purple, FliC-FlgL = Blue, FliC-FlgK = Turquoise, Multi-protein interaction = Orange. D) Interacting residues mapped onto the S. Typhimurium LT2 amino acid sequence. Coloured as above.

**Table 1.** RING interaction network summary between S. Typhimurium LT2 FliC residues (aggregated from across all FliC subunits in both structures) with the hook filament junction proteins (FlgELK), the filament cap (FliD) and other FliC monomers (9GNZ/9GO6).

|  | **FliC** | **FliD** | **FlgL** | **FlgK** | **FlgE** |
| --- | --- | --- | --- | --- | --- |
| **Number of FliC contacts** | 157 | 59 | 67 | 1 | 0 |
| **Contact types** | H-bond,<br>VDW, π-π<br>stack, Ionic | H-bond,<br>VDW, π-π<br>stack, Ionic | H-bond,<br>VDW,<br>Ionic | H-bond,<br>VDW,<br>Ionic | N/A |
| **N-terminal D0/1 (1-178)** | 89 | 31 | 47 | 1 | 0 |
| **Outer domains (179-400)** | 5 | 2 | 0 | 0 | 0 |
| **C-terminal D0/1 (401-495)** | 63 | 26 | 20 | 0 | 0 |

Overall, there were six FliC residues that could interact with three filament proteins depending on their position within the filament, either FliC, FliD and FlgL (n=5) or FliC, FlgL and FlgK (n=1) (Figure S13). All of these calculated multi-partner interacting residues were maintained in N131, N132 and N134 (Figure S13). N131 had 90.7 % of their FliD, FlgL and FlgK interacting residues conserved or replaced with a residue that performed the same interaction type whereas N132 and N134 had 90.1%. When examining FliC-FliC interactions, a total of 157/495 residues form inter-monomer contacts maintained 91,7 %, 91.1 %and 90.4 % of these contacts in N131, N132 and N134, respectively. N132 contains two predicted disruptive changes, N101A and N142A, which lose 16 inter-monomer contacts alongside a FliD interaction and a FlgL H-bond, respectively. N134 was more similar in its interaction network to the functional N131, particularly its interactions with FliD and FlgL (Figure S14). However, the alternative R468K in N134, although chemically similar would replace the guanidinium group with a single amine, potentially leading to the loss of five H-bonds and π-π stacking. N131 maintains many of the key predicted interacting residues found in the *S*. Typhimurium LT2 FliC that form important contacts with FliD, FlgL, FlgK and other FliC subunits (Figure 5C and D, Figure S14).

## Discussion

The bacterial flagellar filament is vital for bacterial motility and mutations in the flagellin sequence have been shown to effect motility in many organisms (5,19,42–46). Here we used ASR to generate conservative ancestral flagellin candidates to attempt to rescue motility in flagellin-deleted strains. We successfully generated an ASR (N131) which supported bacterial swimming in *S*. Typhimurium TM113 but not in *E. coli* SHU649. This generated insight into sequence, structural and expression model perturbations that can influence the formation of functional flagellar filaments.

Both ASRs N131 and N132 were structurally similar to the FliC and FljB proteins of *E. coli* K-12 (8CXM) and *S*. Typhimurium (6RGV), respectively (7,9,47). N134 had a similar structure to *E. coli* O157:H7 (7SN4) (11) when compared with the flagellins that the two expression systems could utilise. There is a large variation of flagellin outer domain structures and sizes (9) and our and others’ previous work has shown that there are stringent structural characteristics of the flagellin domains that can influence the formation of filaments and subsequent motility (19–21,48). Although no motility was observed for any ASR expressed by *E. coli* SHU649, the N131 candidate was able to produce a motile phenotype when expressed by *S*. Typhimurium TM113. Previous work has shown the exchange of proteins between these two closely related genera can complement the deleted host (49), including FliC and FliD (17). Furthermore, the genomes of *S*. Typhimurium and *E. coli* have been successfully hybridised with large chromosomal replacements which showed no reduction in growth fitness compared to their parent strains (50). Due to the diversity of flagellin sequence and tertiary structure (9) we aimed to test conservative ASR flagellin candidates between the divergence node between *E. coli* K-12 and *S*. Typhimurium.

*S*. Typhimurium contains two flagellin homologues, FliC and FljB, which can both form filaments independently (13,51). In *Caulobacter crescentus*, there are six flagellin copies on the genome all of which can be expressed and form filaments that propel the bacteria (52). This differs from *E. coli*, which only contains one flagellin gene. This study has shown that expression and functionality of an ancestral Enterobacteriaceae flagellin gene was only functional in *S*. Typhimurium.

The motility of the N131 expressing *S*. Typhimurium was significantly reduced compared to wild type FliC. Mutant flagellins have also been shown to reduce motility, even when a single mutation is made to the core domains or whole domains of the flagellin are exchanged (19,21,53,54). However, N131 showed a significant increase in the tumble frequency when compared to wild type. A previous study observed that mutations in the flagellin genes of *Rhizobium meliloti* lead to an increasing tumbling phenotype (55). Conversely, when passaged over nine swim plates, although there was an increase in swim ring diameter, there was no change in swim speed for N131.10 variant compared to the original N131.1. Furthermore, analysis of the cells indicated that tumbling frequency significantly increased compared to the parental N131.1. This counterintuitive result may arise from alterations in filament bundling due to the mutation M144I which provides a more similar contact between other FliC and FlgL subunits to that of *S*. Typhimurium FliC/FljB. This increase in tumbling may allow the N131.10 mutant to be optimised for motility in viscous environments as demonstrated by Licata *et al*., who demonstrated that increased tumbling may aid in the navigation of more complex environments (56).

Analysis of the interaction network of the *S*. Typhimurium LT2 FliC with FliD, FlgL, FlgK and FlgE from the cryo-EM structures published by Einenkel *et al* (41), showed that FliC has the majority of its key interacting residues in the core D0/1 domains. When compared with N131, N132 and N134, we found that there was poor residue conservation in the hypervariable outer domains, yet there are conserved interacting residues in the core domains. However, both N132 and N134 had amino acid alterations that would severely impact the atomic bonds that maintain filament stability, such as R468K. This demonstrates that although there is a level of tolerance to the formation of the flagellar filament in *Salmonella*, even minimal changes to the flagellin amino acid sequence are enough to prohibit filament formation, which has been demonstrated in many investigations (57,58,52,59–61). This further shows that, although neighbours on the phylogenetic tree of life, the flagellin export and filament proliferation dynamics differ between *E. coli* and *S*. Typhimurium.

This study has shown that ASR of flagellin is possible in producing filaments capable of driving bacterial motility in *S*. Typhimurium but not *E. coli*. Although limited insight can be made about the evolution of flagellin as a whole from this work due to the conservative approach use to generate the ancestral flagellin candidates, the conclusions of this study do illuminate the complexity of filament formation. We have produced a novel hybrid flagellin that allows us to understand the important residues and structural characteristics required for filament formation and subsequent motility. This work expands our understanding of the intricate requirements needed in the production of the flagellar filament.

## Acknowledgements

J.S: Conceptualisation, investigation, formal analysis, validation, visualisation, writing – original draft, writing - review and editing. J.S.P: Software, validation, writing - review and editing. P.R: Investigation, resources, writing - review and editing. T.M: Resources, writing - review and editing. Y.S Resources, writing - review and editing. M.A.B.B: Conceptualisation, supervision, funding acquisition, writing – original draft, writing - review and editing.

M.A.B.B is supported by the by the Australian Research Council Discovery Project DP240100462. Y.S is supported by JSPS KAKENHI Grant Number JP26K09246.

The authors would like to express our thanks to Prof. Tohru Minamino for the kind provision of the *S*. Typhimurium TM113 strain.

## Ethics Statement

No human or animal subjects or materials were used in this study.

## Data Availability Statement

All the data used are presented as tables and figures in the Supplementary Information.

---

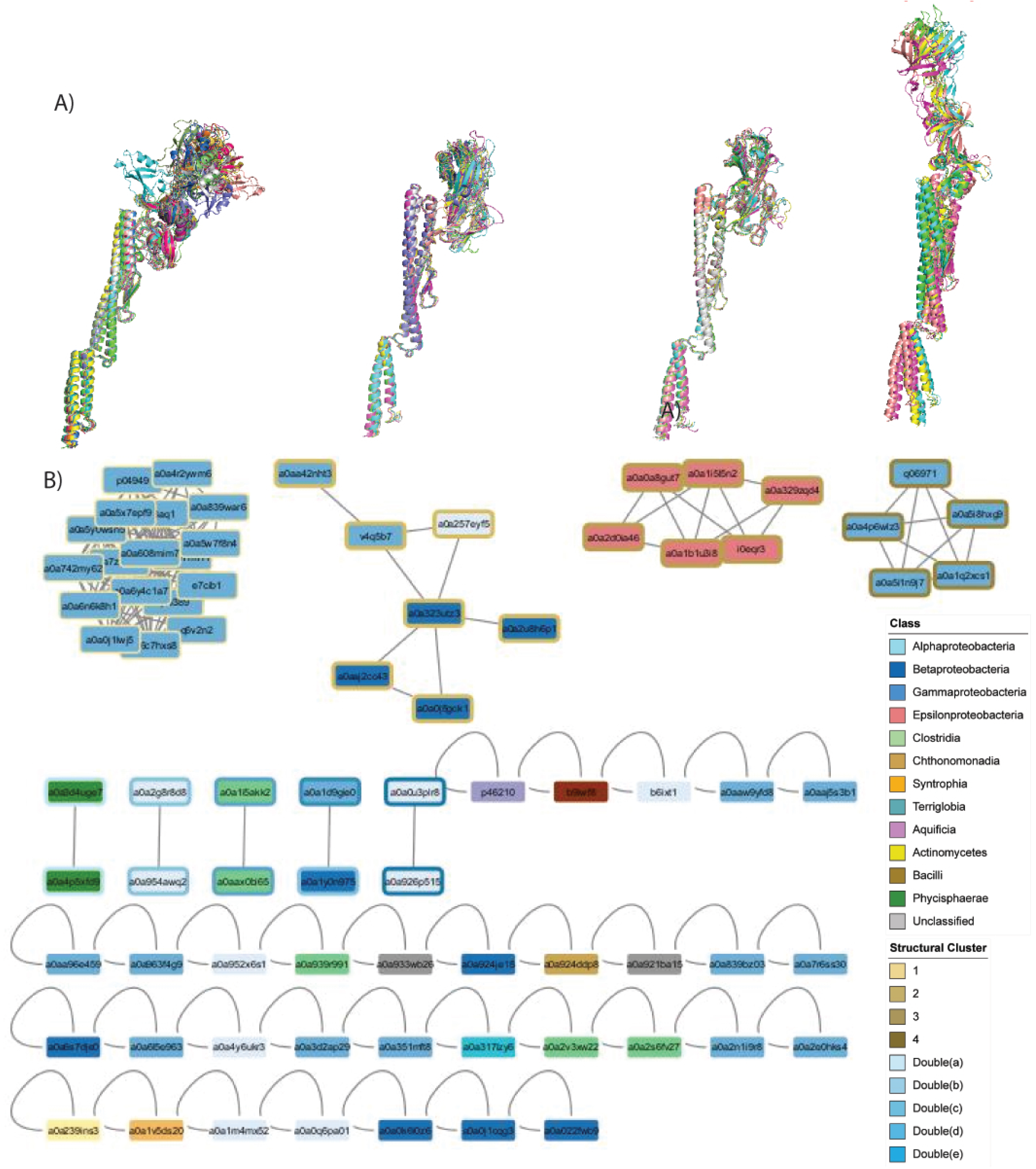

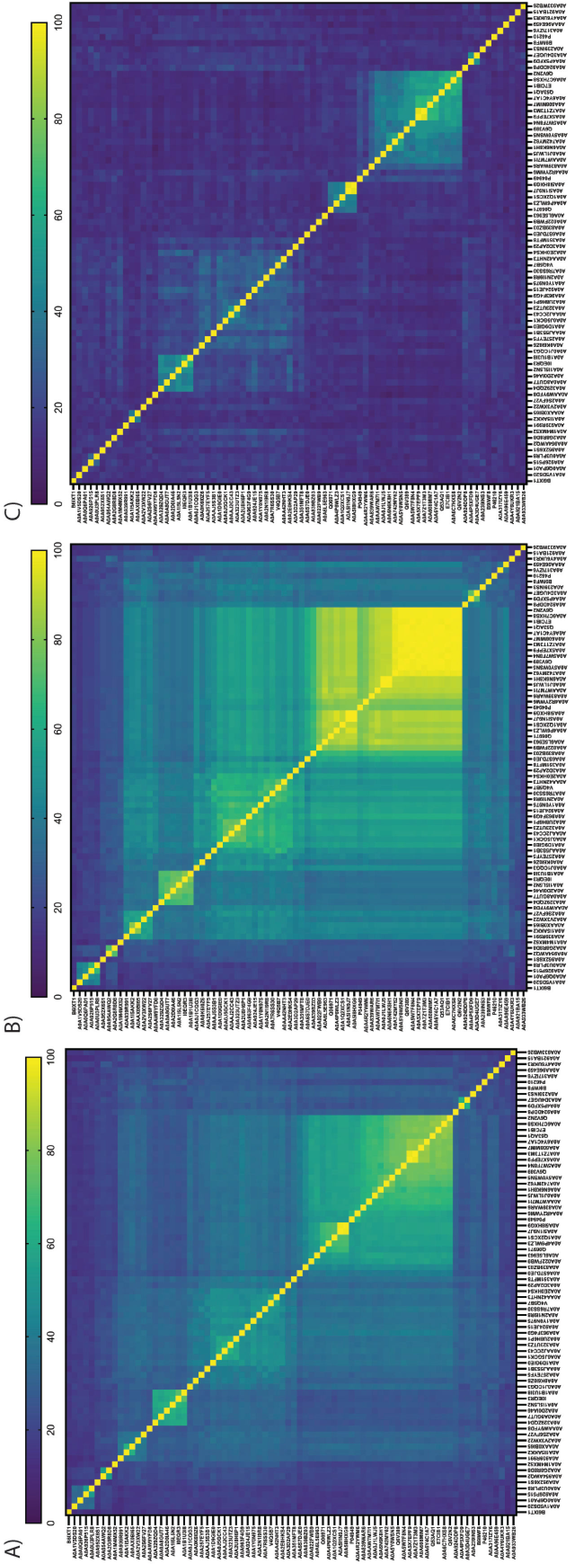

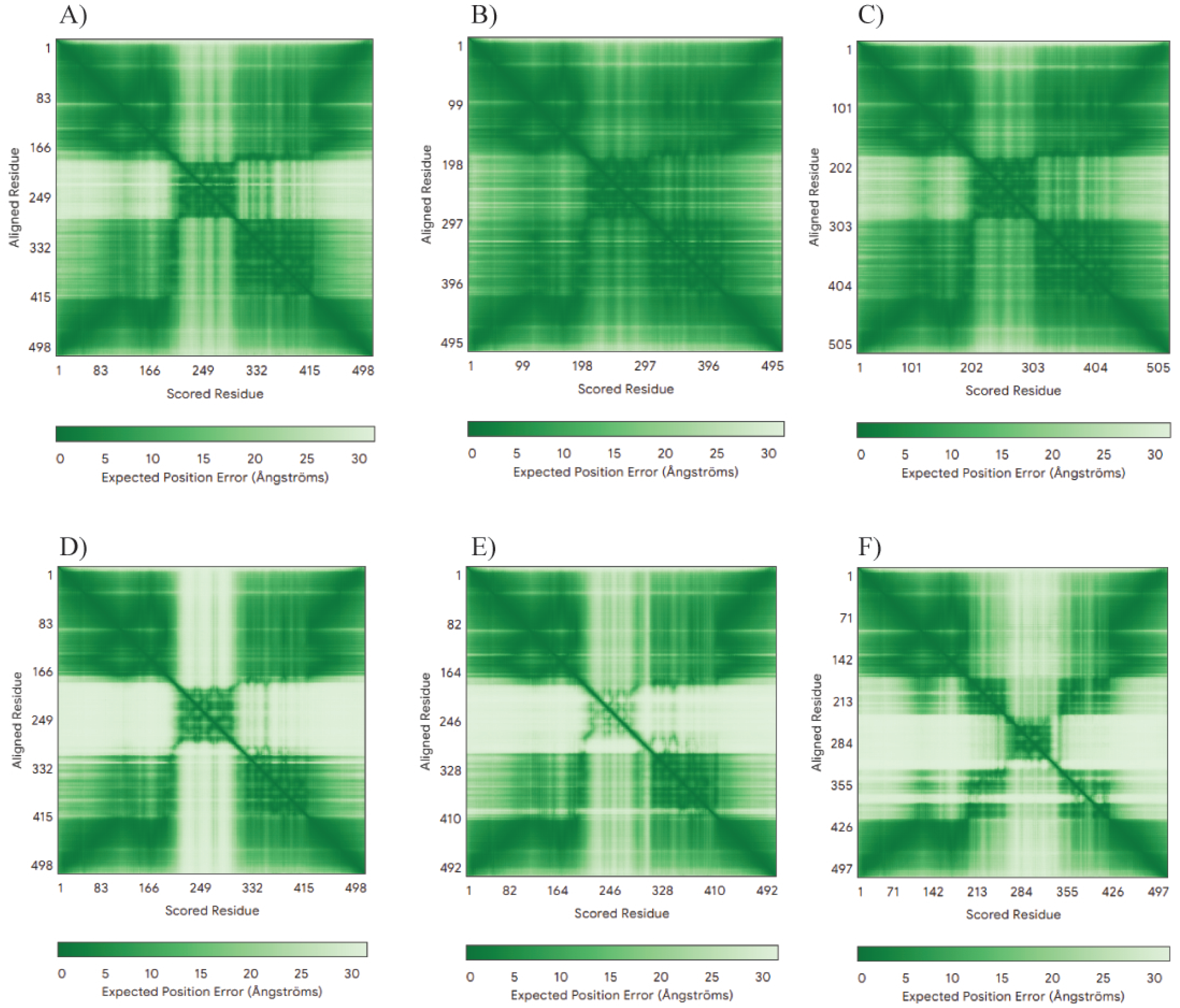

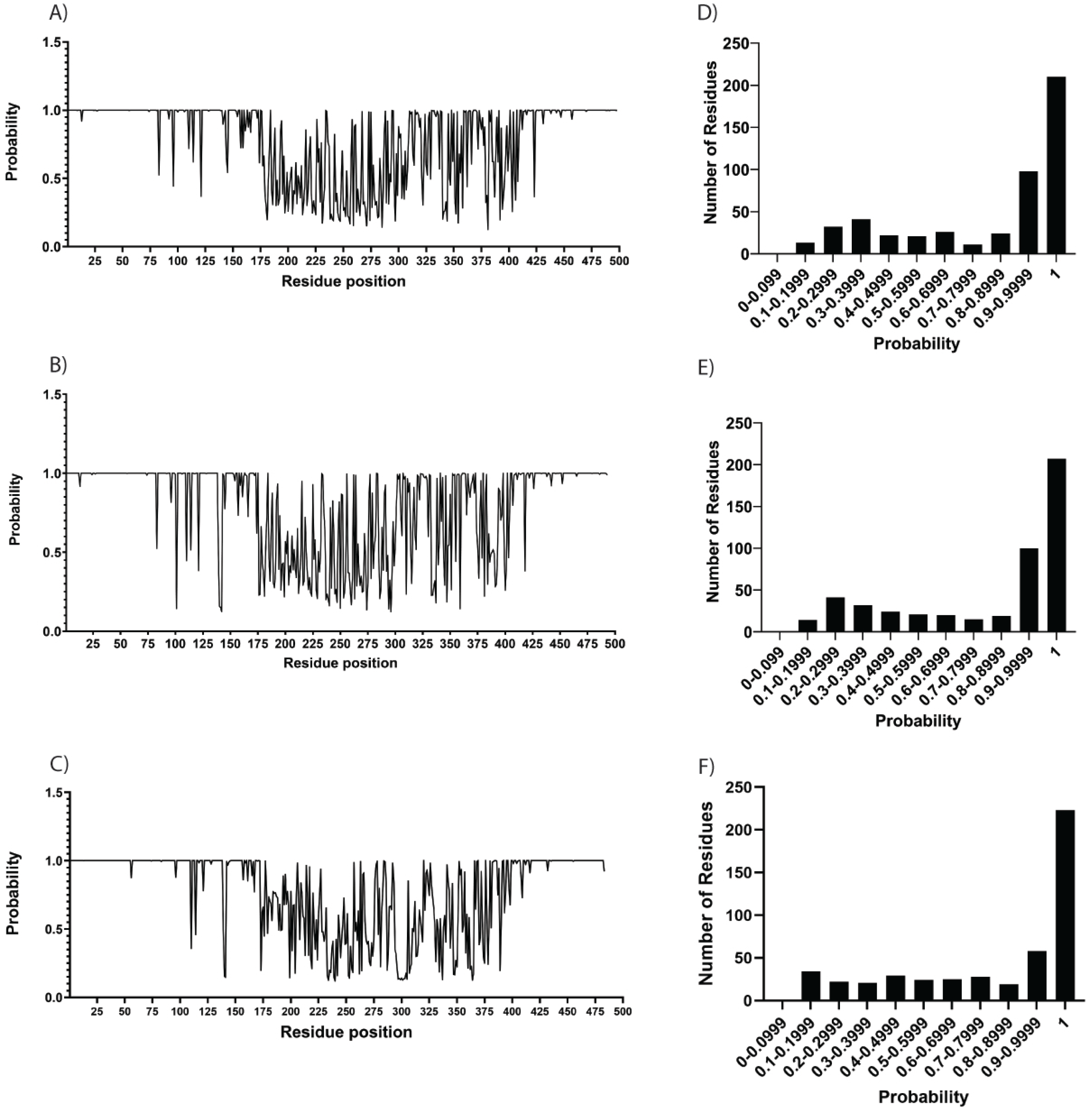

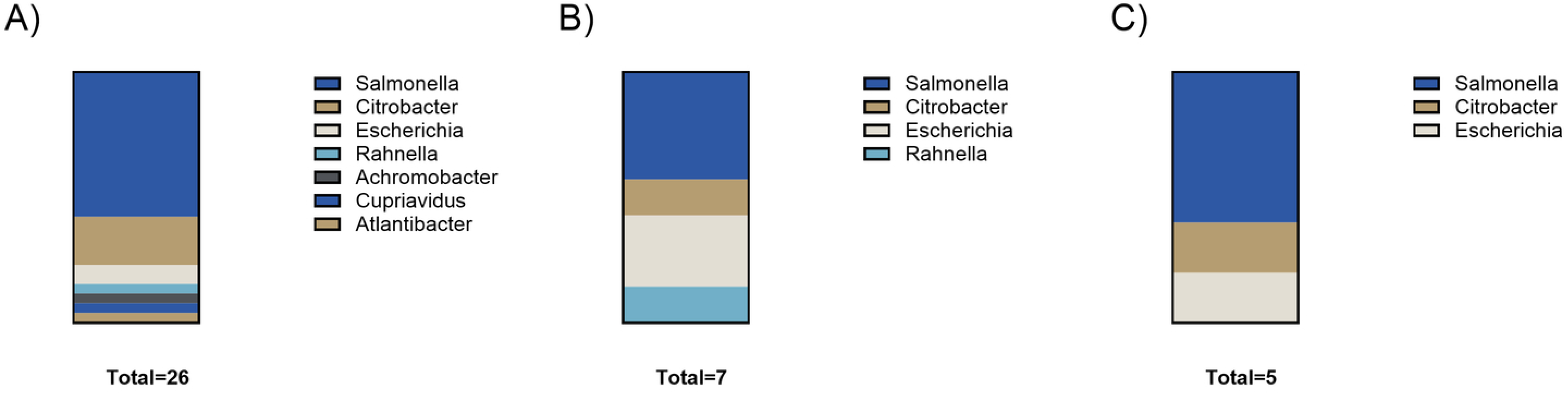

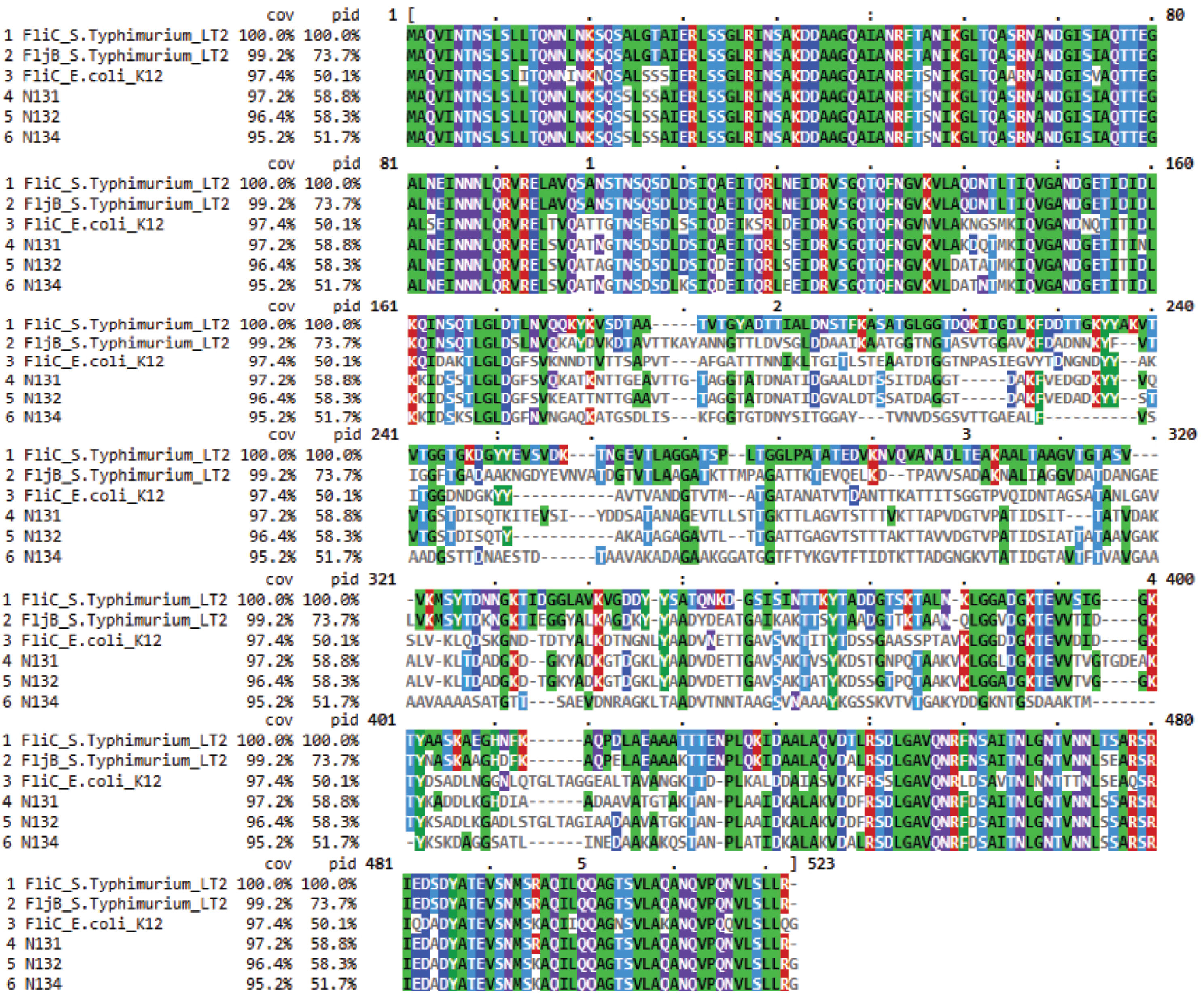

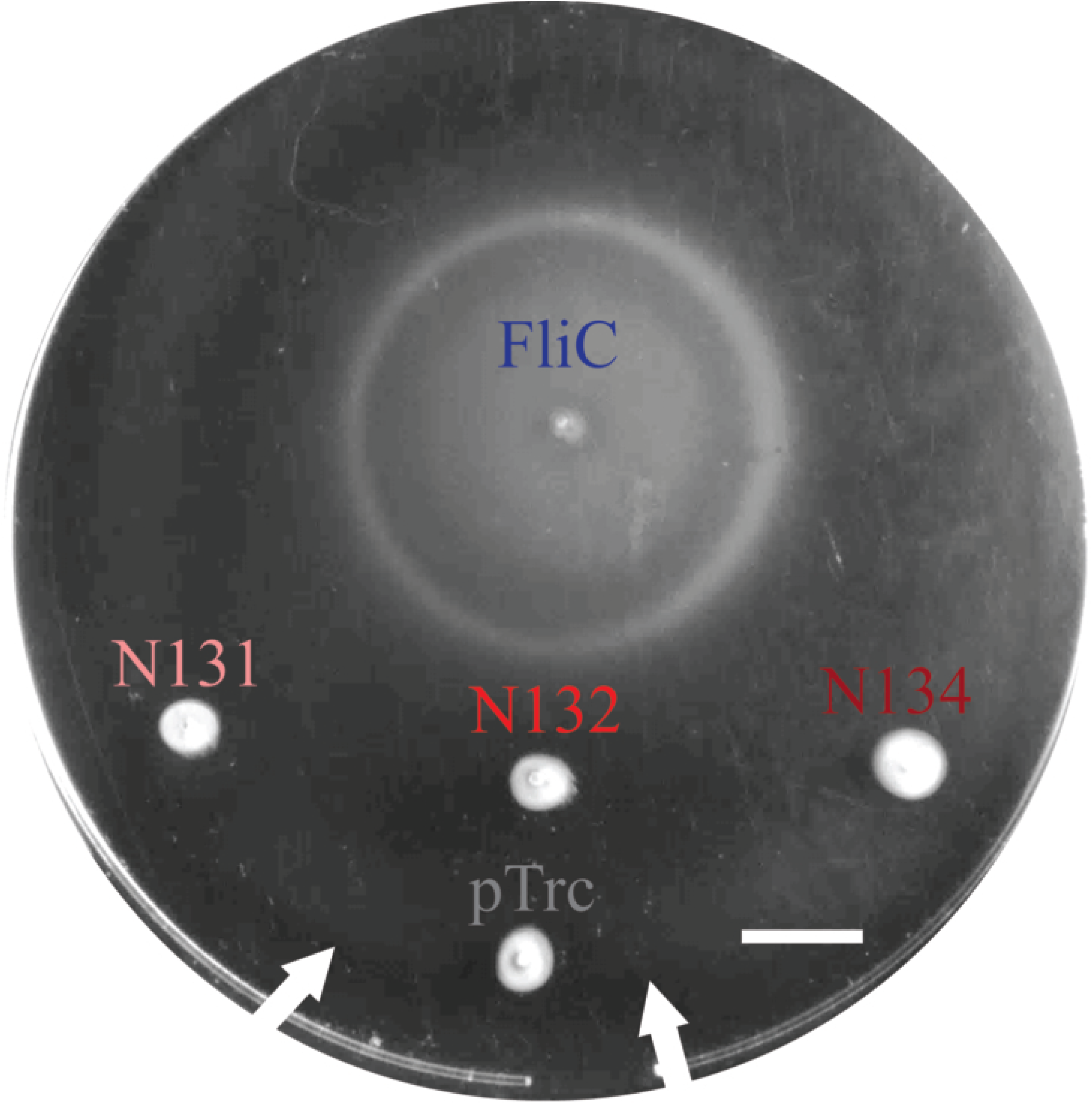

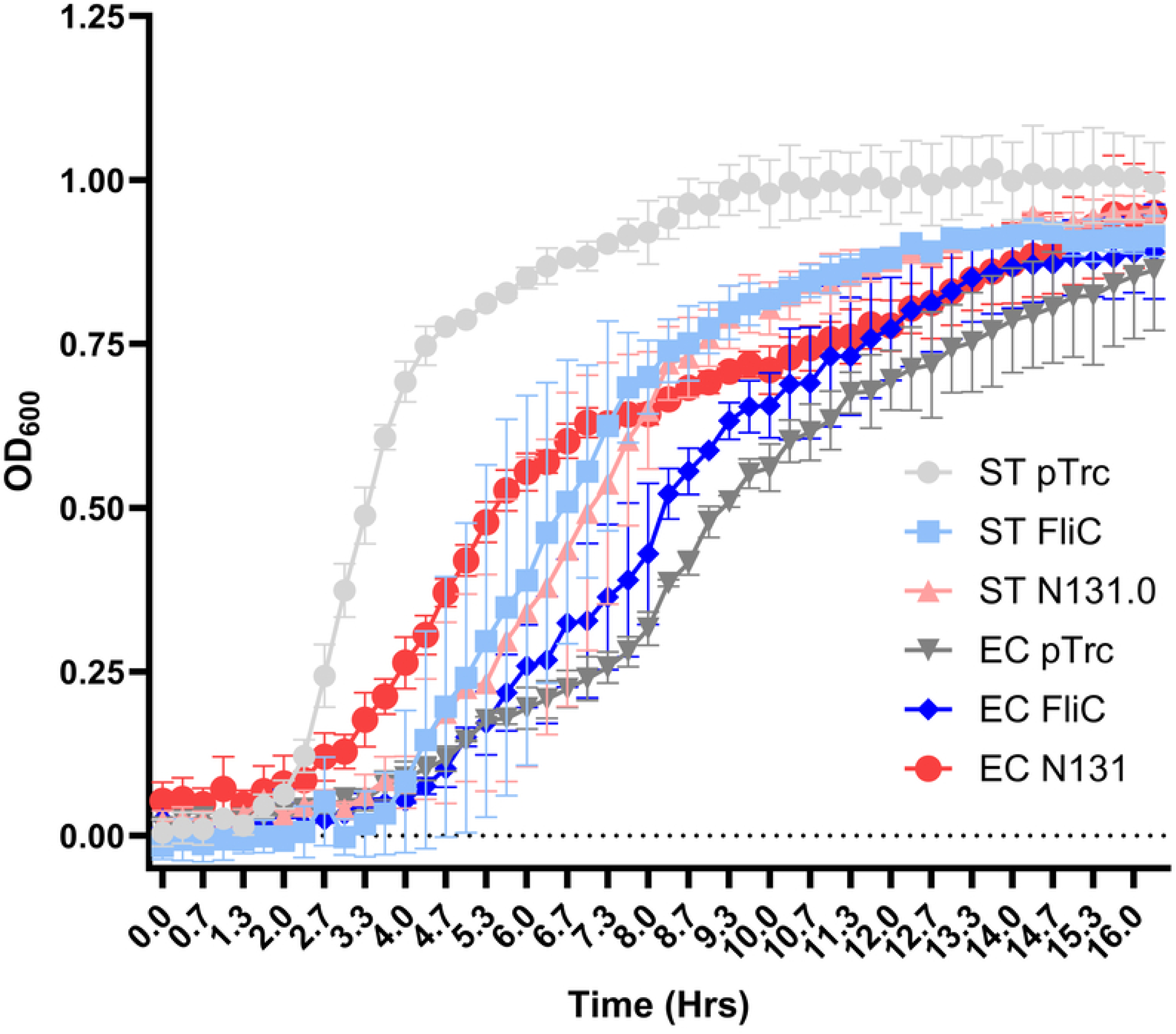

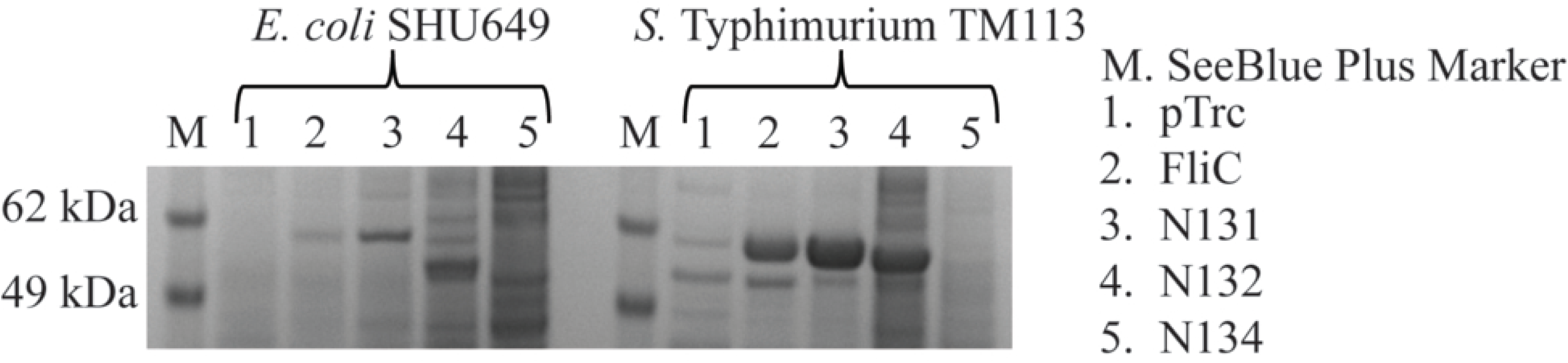

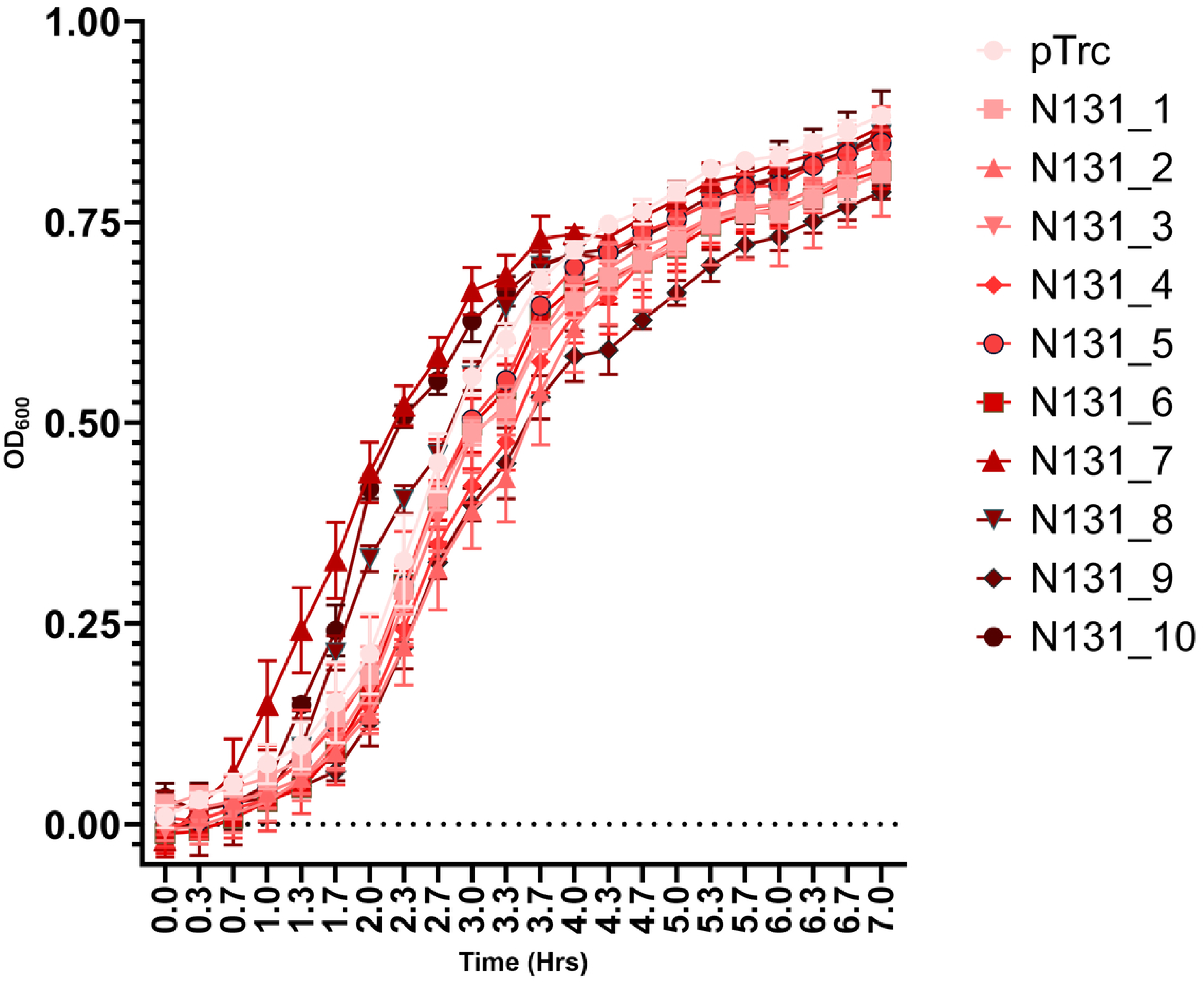

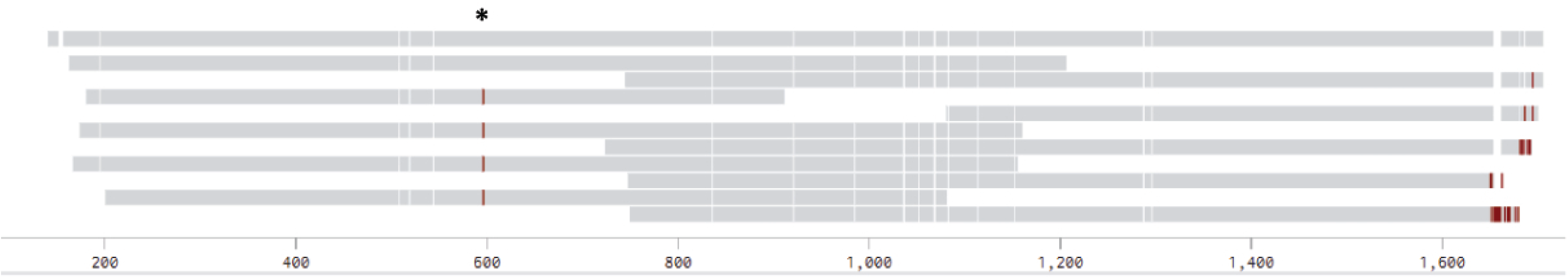

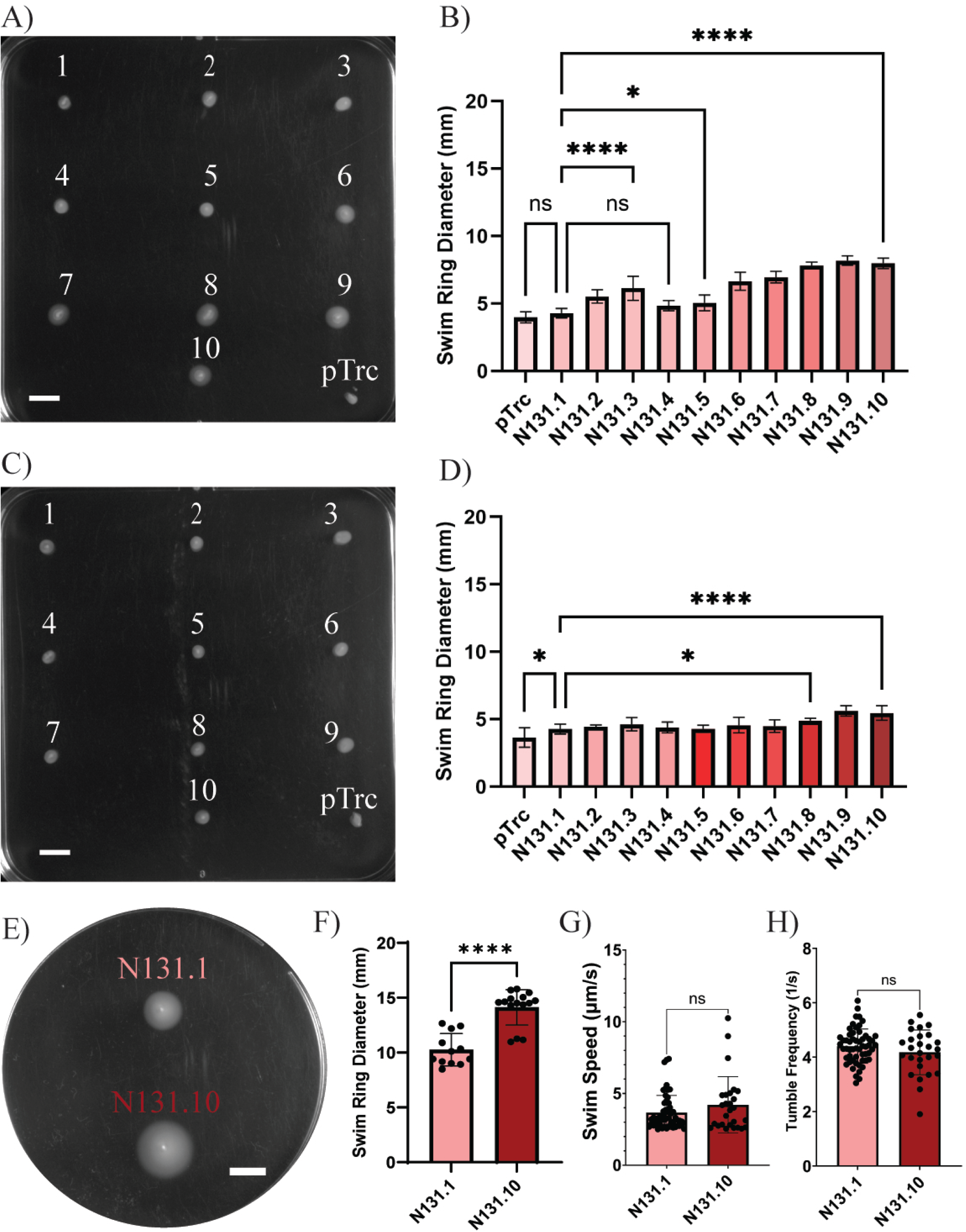

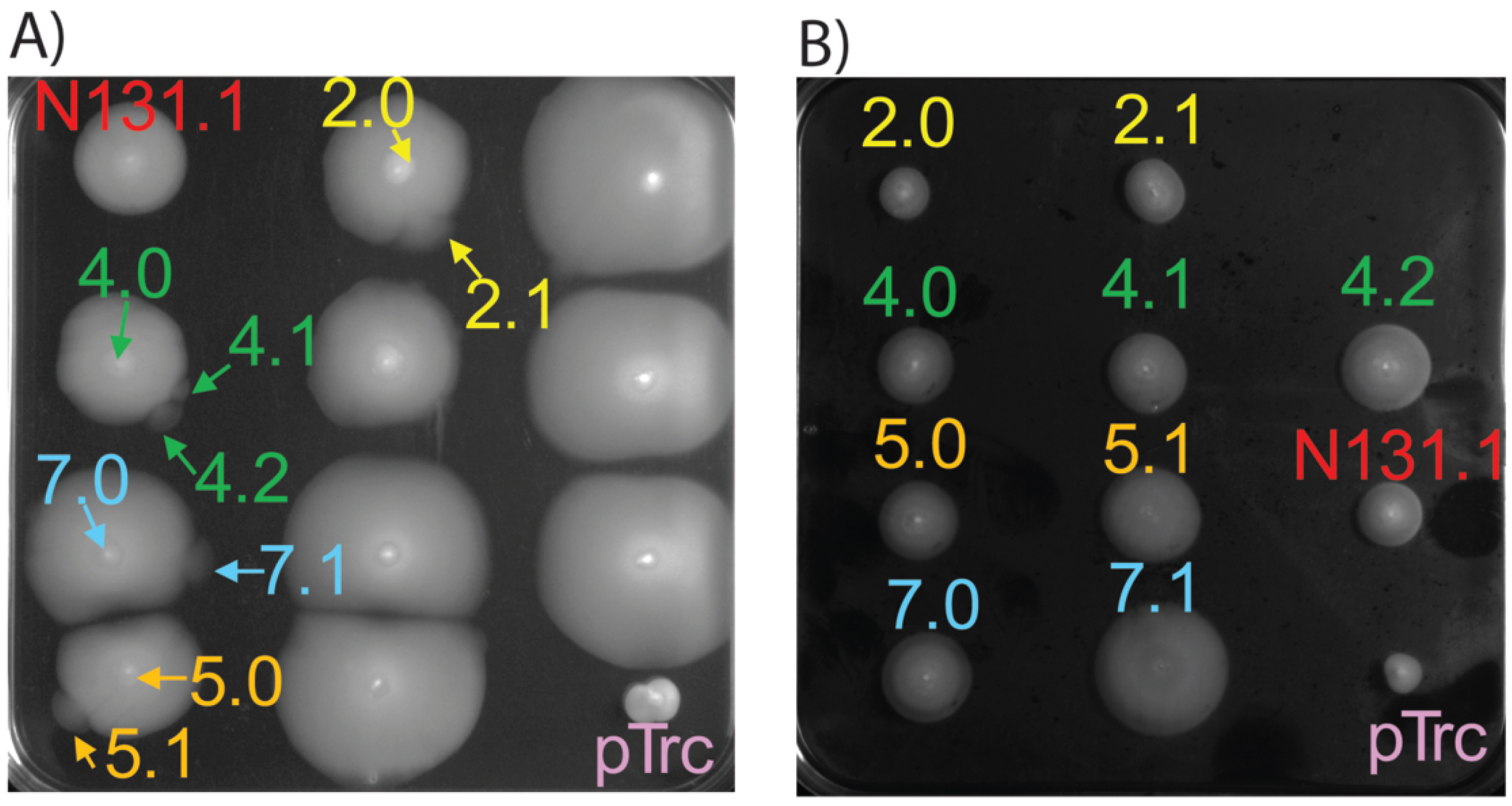

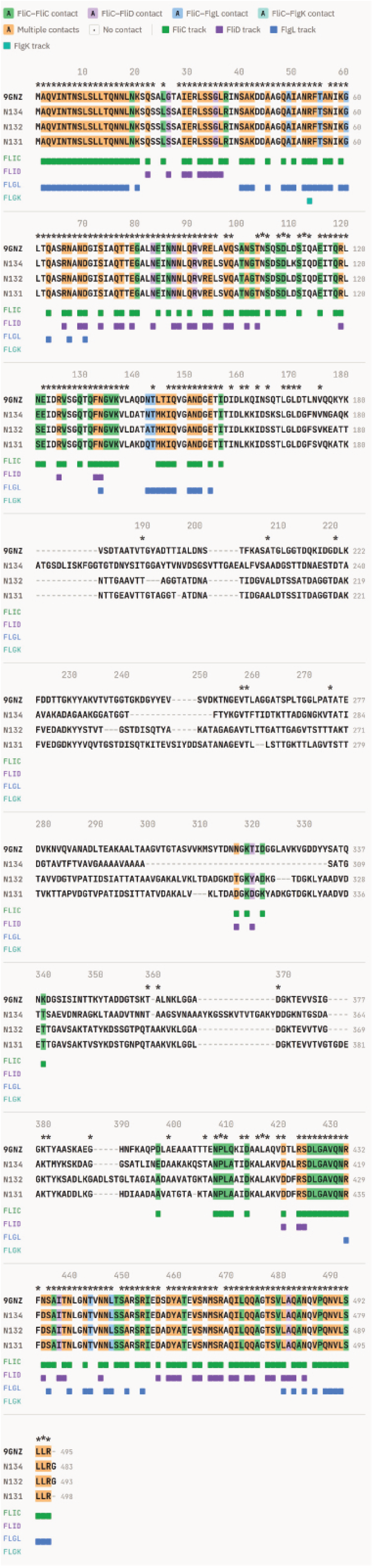

